# Distributional shifts change the biodiversity–ecosystem stability relationship under climate change

**DOI:** 10.1101/2024.01.31.578304

**Authors:** Yuki Kanamori, Shota Nishijima, Ryo Misawa, Takashi Seto, Yoji Narimatsu

## Abstract

Distributional shifts under climate change are increasingly recognized as a biotic change worldwide. However, the effects of distributional shifts on ecosystem variability through changes in biodiversity remain to be clarified. In this study, to elucidate the impact of population-level distributional shifts under climate change on community structure and ecosystem function through ecological hierarchy, we investigated the temporal changes in alpha and beta diversity, the contribution of individual species to the observed changes in biodiversity, and the changes in the relationship between biodiversity and ecosystem variability, using over 25 years of data on about 700 species. We found that ecosystem variability was largely stabilized by beta diversity, especially in the present period (2012 to 2021), because beta diversity increased due to the invasion of species that were distributed mainly in tropic and subtropic areas. Despite the effect of beta diversity, ecosystem variability increased in the present period. This increase in ecosystem variability was caused by the spatially synchronized temporal variation in sea-bottom temperature, which resulted in synchronized temporal variation in species abundance among local communities, as well as the increased impact of alpha variability on ecosystem variability. These results indicate that the impact of alpha diversity on ecosystem variability is more changeable than the impact of beta diversity on ecosystem variability under climate change, suggesting the importance of focusing on changes in alpha diversity resulting from local colonization from other habitats and local extinction in existing habitats.

## 1 Introduction

Distributional shifts under climate change are increasingly recognized as a biotic change worldwide (Walther et al. 2002, Poloczanska et al. 2013). Previous studies have focused on distributional shifts caused by climate change at the species level and the differences in the shifts among species (Pinsky et al. 2013, Poloczanska et al. 2013). However, it remains to be clarified how the distributional shifts of each species impact community structure (e.g., biodiversity) and ecosystem function (e.g., ecosystem variability) through ecological hierarchy.

Alpha diversity (i.e., the number of species within a local community) and beta diversity (i.e., the spatial differences in species composition among local communities) are measurements of community structure, which can be impacted by distributional shifts under climate change as follows. Given that the rate of range contractions in the tail area is slower than the rate of range expansions in the front area (Poloczanska et al. 2013), species distributed in southern areas (or lower elevations) will expand into northern areas (or higher elevations). This can lead to an increasing number of species (i.e., an increase in alpha diversity) and the spatial homogenization of species composition (i.e., a decrease in beta diversity). Indeed, several studies have reported increases in alpha diversity in northern areas and higher elevations (e.g., Walther et al. 2002, Menéndez et al. 2006, Antão et al. 2020) and decreases in beta diversity (e.g., Dornelas et al. 2014, Magurran et al. 2015).

Ecosystem variability (i.e., temporal variability of regional ecosystem productivity) is an ecosystem function. It is thought to be impacted by changes in alpha and beta diversity due to distributional shifts under climate change because recent theory predicts that both alpha and beta diversity can affect long-term ecosystem variability at larger spatial scales through different pathways (Q3 in Fig. 1, Wang and Loreau 2016). According to the theoretical prediction, ecosystem variability can be partitioned into two components: alpha variability (i.e., temporal variability at the local scale) and spatial asynchrony among local communities. As a famous experiment that plots with more species show less year-to-year variation in total biomass (Tilman et al. 2006), alpha variability relates to alpha diversity although alpha diversity can also affect spatial asynchrony among local communities (Wang and Loreau 2016). Therefore, the increase in alpha diversity due to distributional shifts under climate change is expected to decrease alpha variability, resulting in decreasing ecosystem variability. In contrast, given that communities with a more homogenous species composition are expected to exhibit more synchronous responses to a common environment than those with a greater variety of species (Wang and Loreau 2016), spatial asynchrony among local communities relates to beta diversity. Therefore, the decrease in beta diversity due to distributional shifts under climate change is expected to decrease spatial asynchrony among local communities, resulting in increasing ecosystem variability. A previous study found a relatively stronger effect of alpha diversity on ecosystem variability compared with the effect of beta diversity (Wang et al. 2021). However, it is not known how the relative contributions of alpha and beta diversity to ecosystem variability change under climate change.

**Fig. 1:**
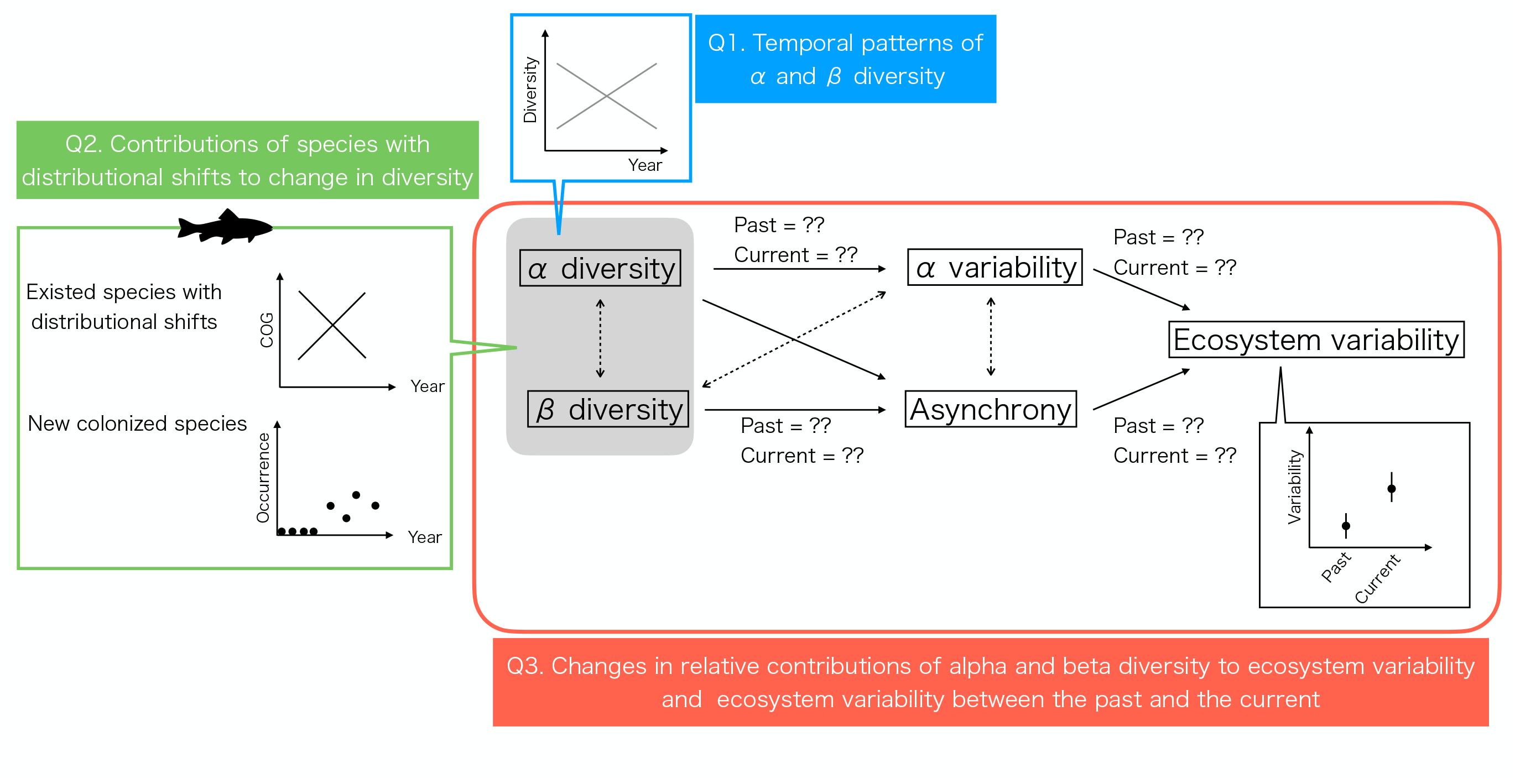
Study design.

The western North Pacific is one of the richest fishing grounds in the world (Takakura 2021), meaning that a wide variety of species are distributed in the area. This is supported by the complex marine environment resulting from three ocean currents: the Oyashio Cold Current, the Tsugaru Warm Current, and the Kuroshio Warm Current. Recently, sea temperature has been increasing (Kakehi et al. 2021), and distributional and phenological shifts have been observed (Kanamori et al. 2019, Kakehi et al. 2021, Kanamori et al. 2023). Scientific surveys have been conducted since the middle 1990s to monitor the bottom species in this area.

In this paper, to elucidate the impact of population-level distributional shifts under climate change on community structure and ecosystem function through ecological hierarchy, we first investigate the temporal patterns of alpha and beta diversity using 25 years of data on bottom species (ca. 700 species and ca. 1500 sites) in the western North Pacific (Q1 in Fig. 1). We then examine the contributions of individual species, especially those that have experienced distributional shifts, to the observed changes in alpha and beta diversity (Q2 in Fig. 1). Finally, we quantify how the relative contributions of alpha and beta diversity to ecosystem variability through alpha variability and spatial asynchrony changed between the past and present periods and how the amount of ecosystem variability changed as the consequence of the change in the relative contributions (Q3 in Fig. 1).

## 2 Materials and Methods

### 2.1 Data sets

The surveys were conducted between 1997 and 2021 in autumn (late September to early November) by the R/V *Wakataka-maru* at depths of 150–900 m along lines A to H (Fig. 2) during the daytime, with a mean ship speed of 3.0 knots for 30 min. The mesh size of the net was 50 mm and a cover net with 8 mm mesh was set at the cod-end. At each station, experts identified species, measured the total weight of each species, and calculated the swept area as survey efforts, using the arrival and departure points on the bottom and horizontal open width of the net recorded by the Net Recorder system (Furuno Electric Co., Hyogo, Japan or Marport, Reykjavik, Iceland).The density data for each species were calculated as the catch weight divided by the survey effort at each station. A total of 680 species were observed across 1,512 sites during the 25-year study period. The top 10 species with the highest average density were *Coelorinchus macrochir,Gadus chalcogrammus,Synaphobranchus kaupii,Bothrocara zestum,Gadus macrocephalus,Etmopterus lucifer,Coryphaenoides pectoralis,Laemonema longipes,Todarodes pacificus*,and *Sebastolobus macrochir*.

**Fig. 2:**
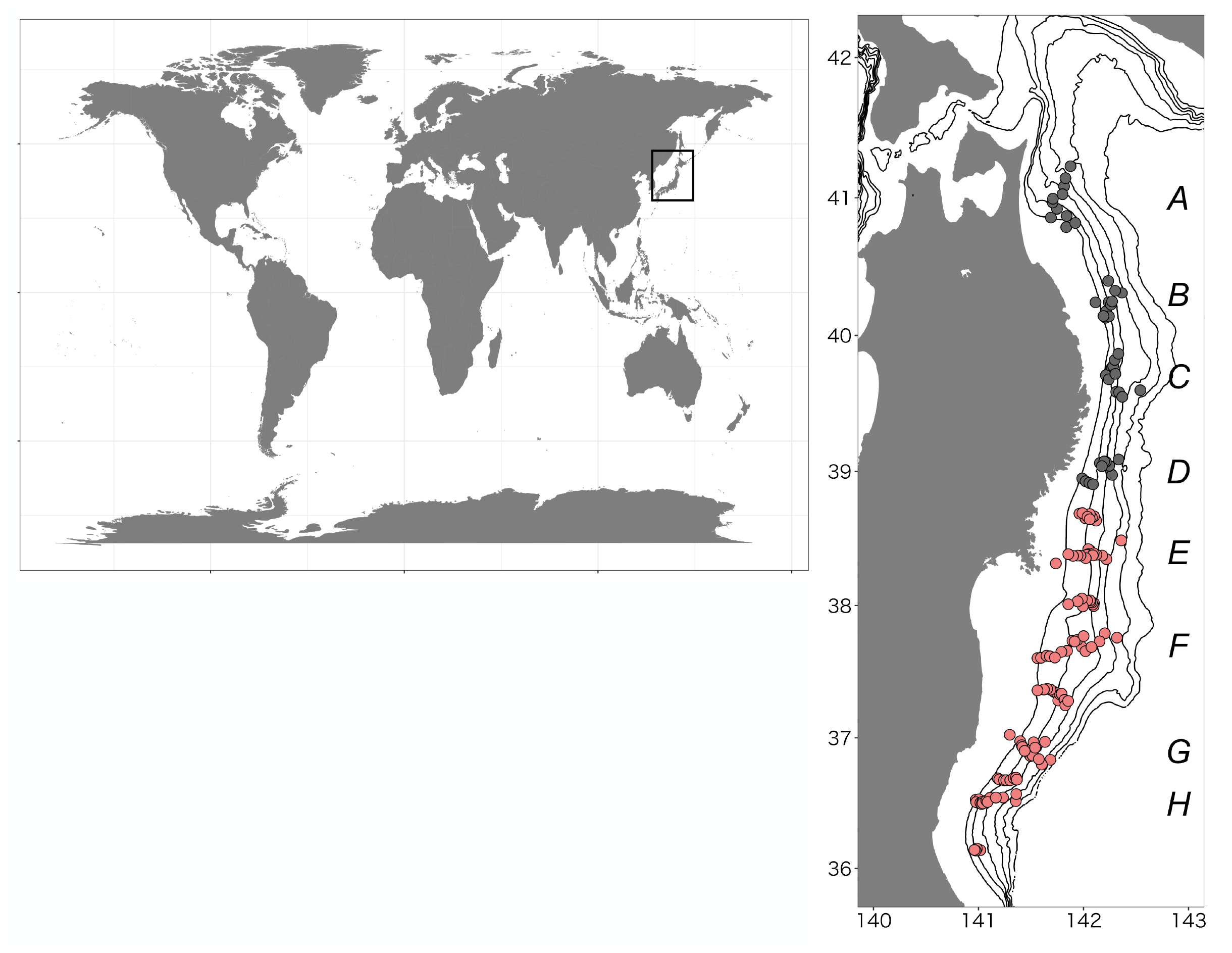
Study area.

Oceanic features in the target region are very complicated. First, there are two fronts, called the Oyashio and the Kuroshio ones where the temperature rapidly changes at the southern edge and the northern edge, respectively. Second, an area, known as the mixed water region, forms between these fronts, where there is not only mixed water originated by both the Oyashio and the Kuroshio, but there are also cold-water eddies with lower water temperatures separated from the Oyashio front, and warm-water eddies with higher water temperatures from the Kuroshio front. In this study, reanalysis of a data assimilation system MOVE (MRI Multivariate Ocean Variational Estimation) of the Meteorological Research Institute of the Japan Meteorological Agency (Fujii and Kamachi 2003; Usui et al. 2006), which highly reproduce the features in our target area (Fujii and Kamachi 2003), has been applied. The data are assimilated using a three-dimensional variational method with satellite SSH, SST which produced by the Japan Meteorological Agency, and water temperature and salinity profiles. The horizontal resolution is 0.1 × 0.1 degrees, lead to appropriately resolve such eddies. Vertical resolution is 54 layers, and by using the coupled vertical EOF mode for water temperature and salinity, appropriate analysis values are reproduced not only for the ocean surface but also for the sub-surface water temperature. The seafloor temperature in the reanalysis has been used in this study. Most of the study sites showed positive trends in the yearly mean sea-bottom temperature (SBT) (Fig. S1).

### 2.2 Data analyses

#### 2.2.1 Biodiversity

We calculated the alpha and beta diversity for each year, using mean species richness and Whittaker’s beta diversity (*β* = *γ/α*) in the study area, respectively. We also calculated alpha and beta diversity in simulated regions for which the study sites were randomly selected without replicates (see 2.2.4 for details) and confirmed that the patterns were similar between the study area and the simulated regions (Figs. S2-3).

#### 2.2.2 Species that experienced distributional shifts

To examine the contributions of individual species with distributional shifts to alpha and beta diversity, we categorized species into four types: (1) range shift species, (2) new colonizing species, (3) range shift & new colonizing species, and (4) species without distributional shifts. (1) range shift species and (2) new colonizing species were considered to be species that experienced distributional shifts in this study area.

#### (1) Range shift species

To identify “range shift species,” we calculated the center of gravity (COG) for each species, using density data *d*(*s, t*) at time *t* in station *s*. The latitudinal and longitudinal COG at time *t*, COG_Lat._(*t*) and COG_Lon._(*t*), were estimated as

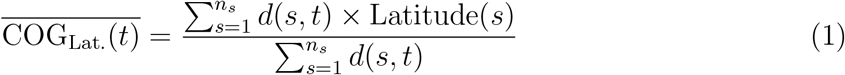

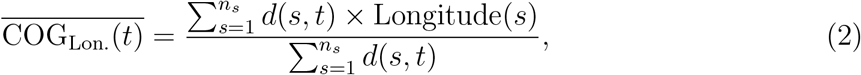

where Latitude(*s*) and Longitude(*s*) are the latitude and longitude of station *s*. We then performed linear regression analyses with COGs as the response variable and year as the explanatory variable for species that had more than 10 data (i.e., species that occurred in the study area more than 10 years). Species for which the linear regression analyses were statistically significant were categorized as “range shift species.”

#### (2) New colonizing species

To identify “new colonizing species,” we examined the temporal occurrence patterns over 25 years. If species appeared in the last 10 years (i.e., since 2012) for the first time, those species were categorized as “new colonizing species.”

#### (3) Range shift & new colonizing species

*Sepia andreana* was the only species that was categorized as both a “range shift species” and a “new colonizing species.”

#### (4) Species without distributional shifts

If species were not categorized as (1) to (3), they were categorized as “species without distributional shifts.”

### 2.2.3 Species that contributed to the changes in alpha and beta diversity

To understand how individual species contributed to the changes in alpha diversity, we determined the main distributions of the new colonizing species, using SeaLifeBase (https://www.sealifebase.ca). Furthermore, we confirmed the spatial patterns where the new colonizing species appeared in the study area for each year.

To understand how individual species contributed to the changes in beta diversity, we calculated the temporal changes in beta diversity for each year (Δ*β*_*total*_) as well as the following four additive components that drove temporal changes in Δ*β*_*total*_ from 1997 to 2021, using the incident-based partitioning method (Tatsumi et al. 2021): decrease in beta diversity via extinctions, Δ*β*_*ext*.*homo*._; increase in beta diversity via extinctions, Δ*β*_*ext*.*diff*._; decrease in beta diversity via colonization, Δ*β*_*col*.*homo*._; and increase in beta diversity via colonization Δ*β*_*col*.*diff*._. Furthermore, each additive component was partitioned into species-level components (Δ*β*^*i*^). In other words, when a regional community has *i* species,

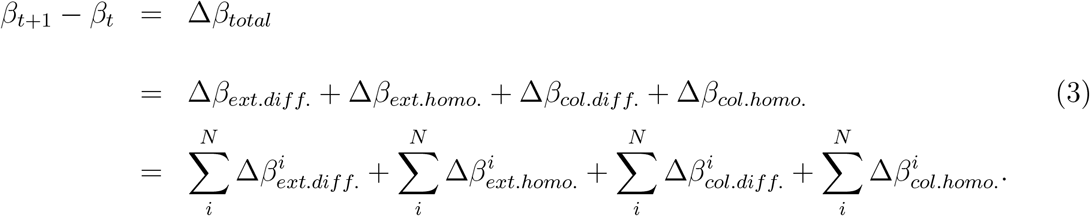

This quantification was performed using the R package *ECOPART* (Tatsumi et al. 2022).

### 2.2.4 Effects of biodiversity on ecosystem variability

To quantify how alpha and beta diversity have affected ecosystem variability through alpha variability and spatial asynchrony, we constructed structural equation models (SEMs) for the data from 1997 to 2010 (past) and from 2012 to 2021 (present) based on a recent study (Wang et al. 2021). We removed the data for 2011 from the analysis because a huge tsunami hit the study area in March 2011 after to the Great East Japan Earthquake and we wanted to exclude the direct effect. The model allowed for ecosystem variability to be partitioned into two components: temporal variability at the local scale (alpha variability) and spatial asynchrony among local communities. In addition, the model included direct paths from alpha diversity to alpha variability and from beta diversity to spatial asynchrony. We also included a direct path from alpha diversity to spatial asynchrony, although the direction of this path was predicted to be context-dependent on the between-site environmental correlation, the between-species environmental correlation, and the strength of the interaction between site- and species-specific environmental responses (Wang and Loreau 2016). Moreover, to account for the effects of unobserved factors, we added correlation errors between alpha diversity and beta diversity, alpha variability and beta diversity, and alpha variability and spatial synchrony.

The components of the SEMs were calculated as follows. Alpha diversity was defined as the mean local species richness (see 2.2.1), and beta diversity was calculated as the ratio of gamma diversity and alpha diversity, where gamma diversity is the regional species richness (see 2.2.1). Spatial asynchrony was defined as 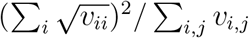, where *v*_*i,j*_ is the temporal covariance in density between site *i* and *j*.

For a direct path from alpha diversity to spatial asynchrony, the changes in between-site environmental correlations were examined. In the between-site environmental correlation, correlations of SBT at site *s*_0_ with respect to SBT at another site *s*_0_ were calculated for the data from 1997 to 2010 and from 2012 to 2021 (e.g., the correlation of SBT for 1997 to 2010 at site *s*_0_ with respect to the SBT for 1997 to 2010 at site *s*_1_). In addition, the differences in SBT at site *s* and site *s* + 1 for each year were calculated using the Bray–Curtis dissimilarity index (i.e., the dissimilarity in SBT between site *s* and site *s* + 1 for year *t*).

Those components of the SEMs were calculated using “simulated regions,” as in previous studies (e.g., Hautier et al. 2018, Ebeling et al. 2020, Wang et al. 2021), because we get only one value of ecosystem stability from one time-series dataset (i.e., the row data of the study area). Simulated regions were made by pooling together *M* plots that were randomly selected from survey sites. We set *M* as 60% and 80% of the total study sites and repeated random selection until 10,000 simulated regions were obtained. We confirmed that the patterns of alpha and beta diversity from the simulated regions were similar to the ones from the study sites (Fig. 3, Figs. S2-3).

**Fig. 3:**
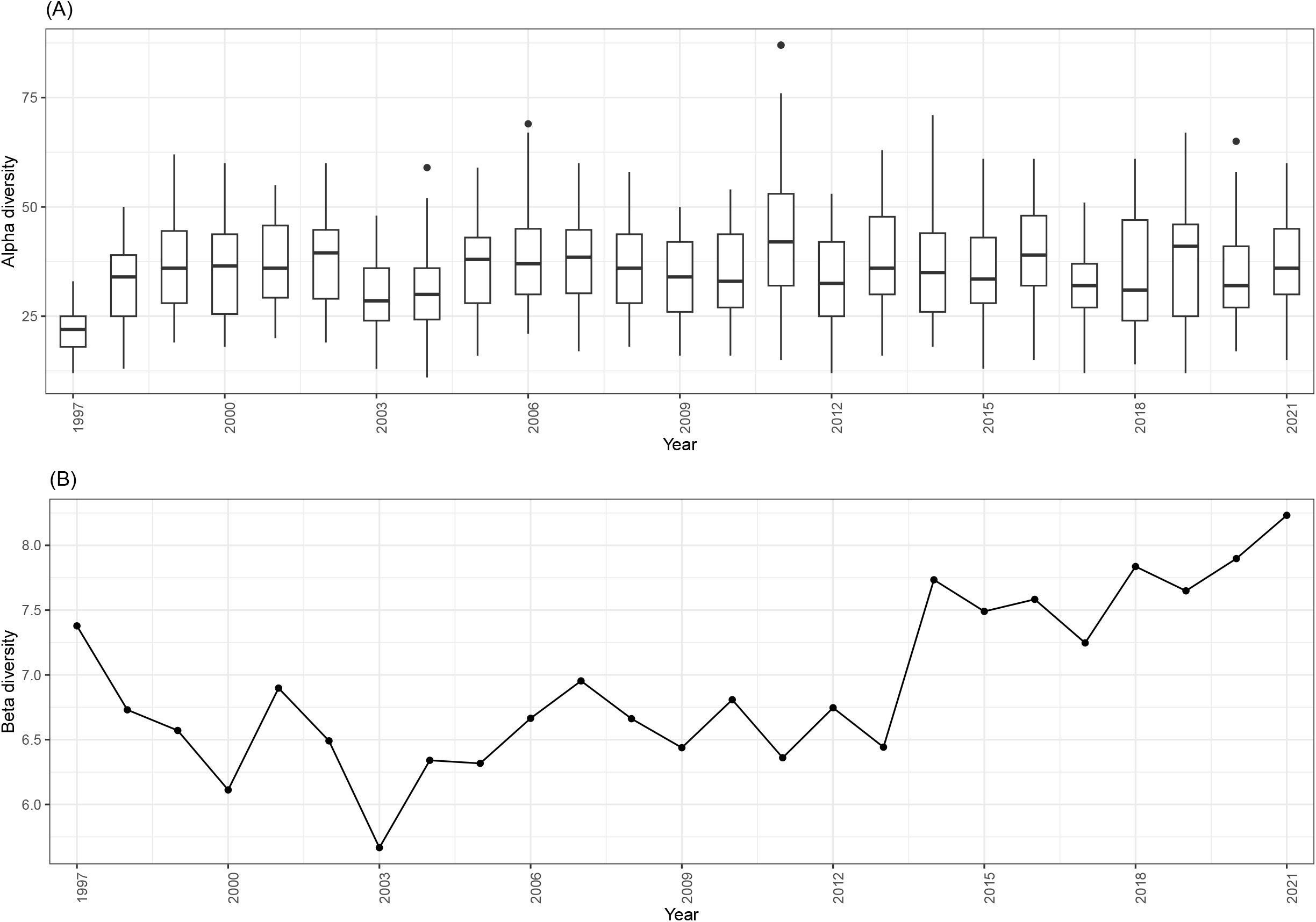
Temporal changes in alpha and beta diversity. Alpha diversity was defined as mean species richness. Beta diversity was calculated using Whittaker’s beta diversity.

SEMs were created using the R package *lavaan* (Rosseel 2012). We used two common index methods to evaluate the goodness-of-fit of the SEMs, namely, standardized root mean square residual (*<* 0.05) and goodness-of-fit index (≧ 0.9), as recommended (Hoyle 2012), because the chi-square test is considered unreliable due to excessive Type I errors for data larger than 200 cases (Kenny et al. 2015).

## 3 Results

### 3.1 Temporal changes in alpha and beta diversity

Alpha diversity did not show a major increasing or decreasing trend, whereas beta diversity showed an increasing trend after 2003 (Fig. 2). The magnitudes of the changes in alpha and beta diversity between 2010 and 2011 were not so large compared with other years, although huge tsunami waves hit the study area in 2011 after the Great East Japan Earthquake.

A total of 133 species newly appeared after 2012, and those species are considered to be new colonizing species in this study. Of these, 43.6% were tropical and subtropical species, 15.0% tropical to temperate species, 23.3% temperate species, and 18.0% species with unknown distribution. Moreover, the number of new colonizing species tended to be higher in the southern part of the study area (Fig. S4).

During the 25-year study period, 72 species showed significant trends in COG (Figs. S5-8), and those species were considered to be range shift species in this study. A total of 23 species, including *Hippoglossoides dubius, Clidoderma asperrimum*, and *Antimora microlepis*, showed increasing trends in the COG of latitude, while 23 species, including *Sebastolobus macrochir, Gadus chalcogrammus*, and *Ommastrephes bartramii*, showed increasing trends in the COG of longitude. In contrast, 20 species, including *Cleisthenes pinetorum, Pseudopleuronectes herzensteini*, and *Pandalus eous*, showed decreasing trends in the COG of latitude, while 21 species, including *Hippoglossoides dubius, Dexistes rikuzenius*, and *Pandalus nipponensis*, showed decreasing trends in the COG of longitude.

Species with distributional shifts contributed to the temporal changes in beta diversity in two ways (Fig. 3). First, range shift species contributed mainly to colonization homogenization and extinction heterogenization at nearly equal rates (30.4% and 30.8%, respectively). In addition, these contributions increased slightly over time. Second, new colonizing species (i.e., species that occurred in recent 10 years) contributed mainly to colonization differentiation and extinction homogenization at rates of 18.0% and 15.2%, respectively.

### 3.2 Effect of alpha and beta diversity on ecosystem variability

The effects of beta diversity on ecosystem variability through asynchrony were similar between the past (1997 to 2010) and the present (2012 to 2021) periods (Fig. 4A,B). In contrast, the effect of alpha diversity and alpha variability on ecosystem variability through asynchrony increased from the past to the present period, leading to an increase in ecosystem variability (Fig. 4C). These tendencies remained when the simulated region was created using 60% of the study sites (Fig. S9).

**Fig. 4:**
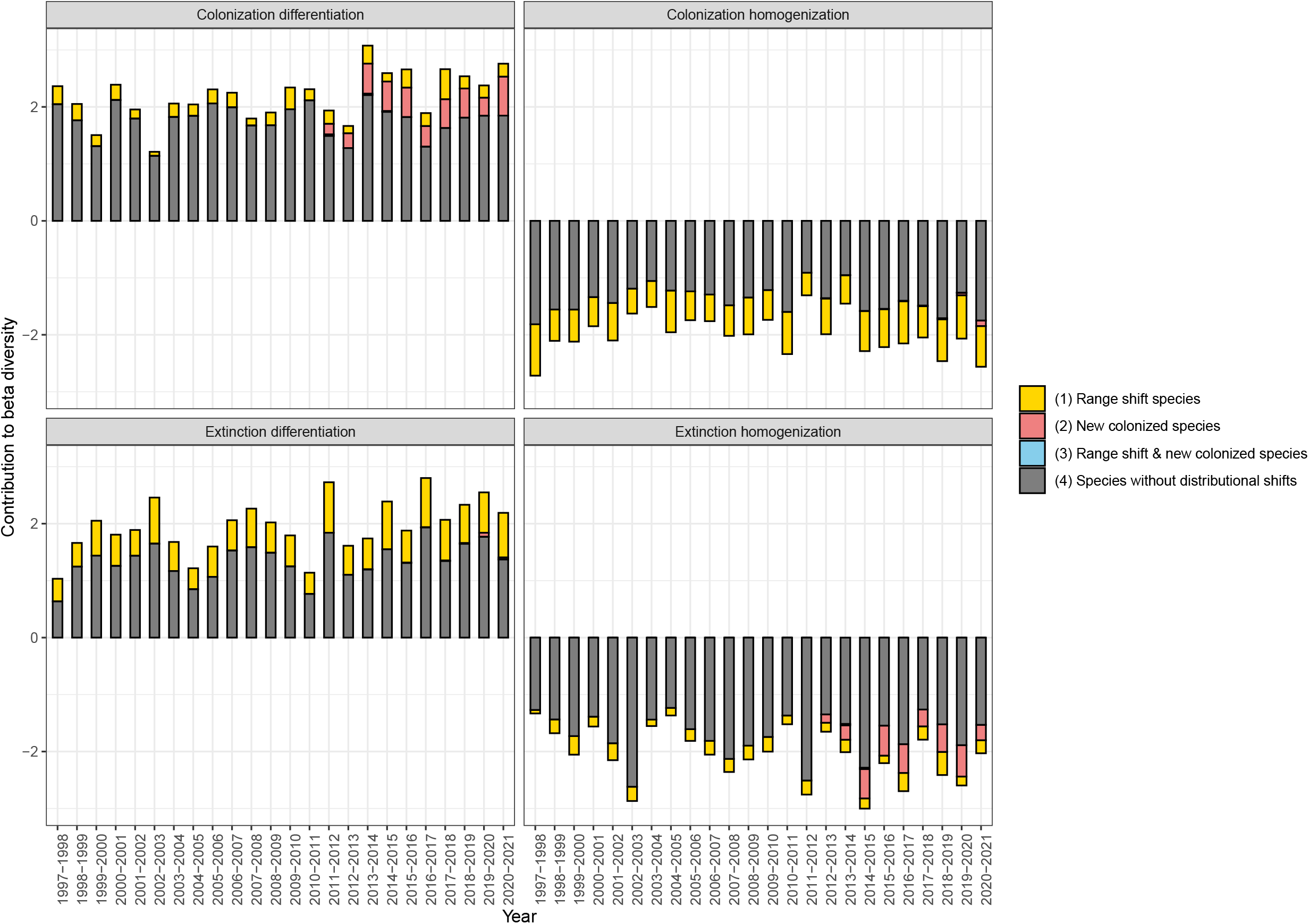
Contributions of individual species to temporal changes in beta diversity.

**Fig. 5:**
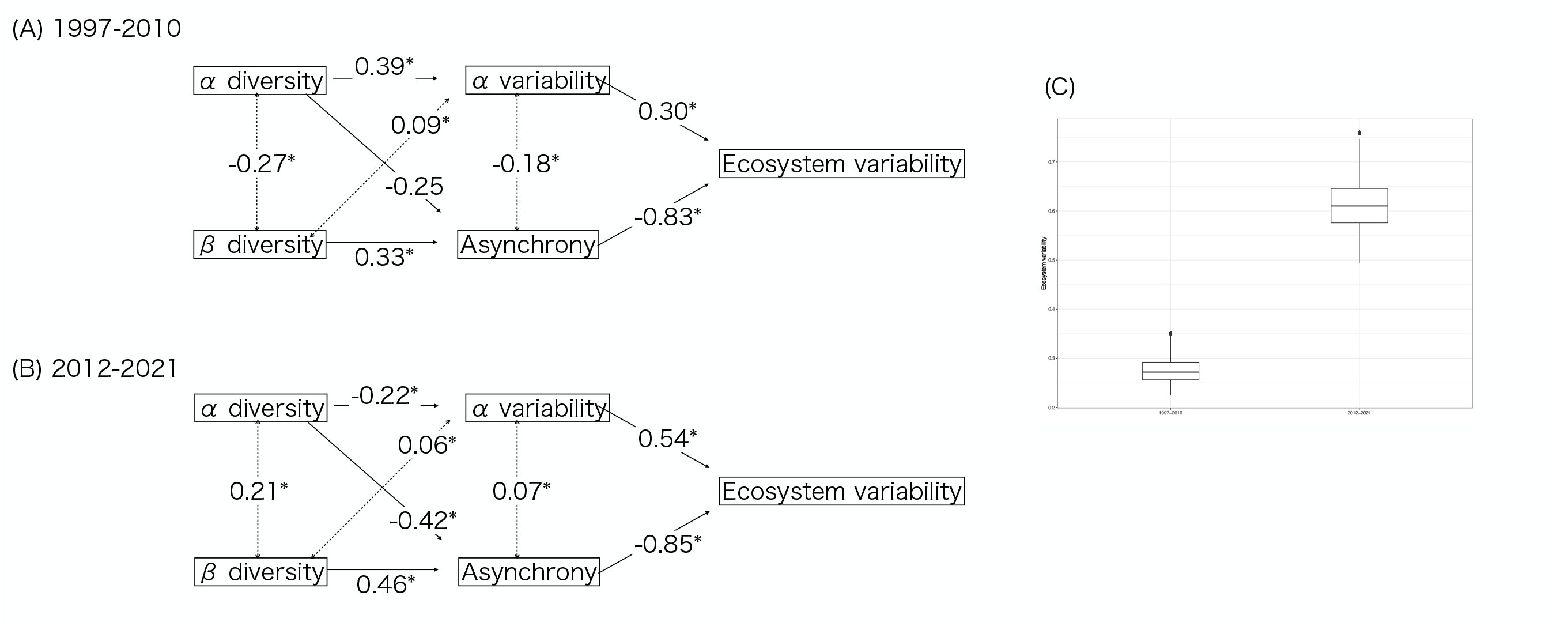
Effects of biodiversity on ecosystem stability in 1997–2010 (A) and in 2012–2021 (B), and ecosystem variability in 1997–2010 (past) and 2012–2021 (present) (C).

For a direct path from alpha diversity to spatial asynchrony, the mean value of correlations of SBT among sites in the simulated region was higher in the present period (0.44) compared with the mean value in the past period (0.24). In addition, the differences in SBT among all study sites were similar over the 25-year study period (Fig. S10).

## 4 Discussion

Distributional shifts under climate change are a well-known biotic change worldwide (Walther et al. 2002, Poloczanska et al. 2013), although the effects on ecosystem function, including ecosystem variability, through changes in biodiversity remain to be clarified. In this study, we investigated temporal changes in alpha and beta diversity, the species that contributed to the observed changes in biodiversity, and the changes in the relationship between biodiversity and ecosystem variability, using over 25 years of data on about 700 species. We found that ecosystem variability was largely stabilized by beta diversity, especially in the present period (2012 to 2021) because beta diversity increased due to the invasion of species that were distributed mainly in tropic and subtropic areas. Despite the effect of beta diversity, ecosystem variability increased in the present period. This increase in ecosystem variability was caused by the spatially synchronized temporal variation in SBT, which resulted in synchronized temporal variation in species abundance among local communities, as well as the increased impact of alpha variability on ecosystem variability. These results indicate that the impact of alpha diversity on ecosystem variability is more dynamic compared with the impact of beta diversity on ecosystem variability under climate change, suggesting the importance of focusing on changes in alpha diversity due to local colonization from other habitats and local extinction in existing habitats.

Species colonization from southern habitats and increased species richness have been well documented under climate change (e.g., Menéndez et al. 2006). In the present study, alpha diversity did not increase, although many species newly colonized the study area.

This may be due to the fact that the colonized new species have not yet become established. Indeed, the temporal occurrence rates of new colonizing species were quite low.

Although decreases in beta diversity have been observed worldwide (e.g., Dornelas et al. 2014), beta diversity increased in area examined in the present study. This disparity may be due to two factors. The first possibility is that the presence of spatial variation in environmental conditions prevents a decrease in beta diversity. For example, a previous study examining similar marine species to our study suggested a decrease in beta diversity occurred due to decreased differences in sea-surface temperature between the northern and the southern areas and because species distributed in the southern area colonized the northern area (Magurran et al. 2015). In contrast, our study area was surrounded by the three currents (Oyashio Cold Current, Tsugaru Warm Current, and Kuroshio Warm Current) and had a significant environmental gradient (i.e., depth). Indeed, spatial variation in SBT persisted in the study area. Another possibility is that we observed the very early stage of colonization from the southern area before establishment. Indeed, we confirmed that the spatio-temporal frequencies of the new colonizing species were quite low. The contributions of new colonizing species to colonization homogenization were quite low as well. Ochoa-Ochoa et al. (2012) revealed temporal changes in the trend of beta diversity under climate change: first comes heterogenization, then homogenization. Therefore, it is possible that the study area will experience a change in terms of the temporal trend in beta diversity, which impacted ecosystem variability through asynchrony, suggesting the need for continued monitoring.

## Acknowledgments

This research was financially supported by the Japan Fisheries Research and Education Agency, and the Fisheries Agency, the Ministry of Agriculture, Forestry and Fisheries of Japan.

## Authorship

YK conceived the research idea. RM and YN conducted the survey and organized the survey data. SS organized the SBT data. YK and SN designed the statistical analyses, and YK wrote the programs and performed the analyses. YK and SS wrote the manuscript with input from all co-authors’ comments.

